# Effectiveness and efficiency of temporary fences as mitigation measures for preventing amphibian roadkill

**DOI:** 10.64898/2026.04.14.718167

**Authors:** Le Brishoual Meven, Tanvier Clara, Dehaut Nathan, Miaud Claude, Jumeau Jonathan

## Abstract

Temporary fences are used to prevent amphibians from accessing dangerous areas. These fences can be built out of different materials and field studies have shown that some of them still allow for trespassing. In this study we compared, in controlled conditions and using experimental arenas, the effectiveness and efficiency of the three most commonly used materials for temporary amphibian fences (polyethylene tarp, polyethylene netting, and wire meshing) and of the presence of an overhang, on two amphibian species with distinct modes of locomotion (*Bufo bufo* for walker/short-distance jumper and *Rana dalmatina* for long-distance jumper). The polyethylene tarp fence, and fences equipped with an overhang, were the only designs able to stop every individual for three consecutive trials. Fences that were not smooth nor equipped with an overhang could be crossed at any height by both species. For non-climbable fences (smooth or equipped with an overhang), long-distance jumper species required a 60 cm high fence to be stopped while the walker/short-distance jumper species only required 20 cm. The walker/short-distance jumper individuals interacted 1.7 times more with the polyethylene netting and wire meshing than the polyethylene tarp, this difference is likely due to the tarp being opaque and impossible to climb, deterring the amphibians from interacting with it. The long-distance jumper species was not deterred by the tarp fence and interacted equally with all materials. When considering price, durability and ease of use while maintaining effectiveness, the polyethylene netting fences equipped with an overhang were the most efficient option for long-term use, while polyethylene tarp is the best option for short-term use. These fences should systematically be equipped with a 10 cm overhang at their top to ensure their effectiveness.

## INTRODUCTION

The life cycle of most amphibians requires them to cross roads separating terrestrial habitats and breeding aquatic habitats (Joly 2019, Cayuela et al. 2020). Amphibians are vulnerable to traffic because they move slowly and can display a freezing behaviour in the face of danger (Carr and Fahrig 2001, Hels and Buchwald 2001, Gibbs and Shriver 2005, Mazerolle et al. 2005). Many species also exhibit temporally concentrated migrations, during which thousands of individuals may simultaneously cross roads and suffer high mortality rates, even under low traffic conditions (Gibbs and Shriver 2005, Orłowski et al. 2008, Helldin and Petrovan 2019). Road mortality surveys which account for amphibians place them as the group with the highest mortality counts or rates, to a point that may lead to population decline (Smith and Dodd 2003, Gibbs and Shriver 2005, Glista et al. 2008, Jochimsen et al. 2013, Beaune 2019).

In order to reduce amphibian roadkill, mitigation measures have been implemented along roadsides known to be mortality hotspots (Puky 2003). These measures include traffic and speed reduction, road closures, translocation of breeding habitat, or limiting road access with fences (Puky 2003, Schmidt and Zumbach 2008, Beebee 2013, Jochimsen et al. 2013). The latter prevents amphibians from getting on the asphalt while guiding them towards underpasses or pitfall traps (Schmidt and Zumbach 2008, Beebee 2013). Fences can be permanent or temporary. Permanent fences vary in material with wire netting, polyethylene netting, concrete or metal often being used, with recommended heights between 45 and 60 cm to prevent climbing (Conan 2023, Gould et al. 2024). The main caveat of permanent fences is their high cost and need for vegetation maintenance (Schmidt & Zumbach 2008). Temporary fences are the most frequently used, representing 84% of roadside installations in France (Eggert 2019), and also being used as exclusion fences for construction sites. They are usually installed during the seasonal migrations of amphibians, or construction duration only. Temporary fences mostly consist of polyethylene netting, wire netting, or opaque polyethylene tarp (Arntzen et al. 1995, Iuell et al. 2003, Morand & Carsignol 2019). Permanent fences are associated with underpasses allowing migrating amphibians to cross the roads. On the contrary, temporary fences are usually associated with pitfall traps, requiring the daily intervention of operators to manually transfer amphibians across the road (Puky 2003, Schmidt and Zumbach 2008, Morand & Carsignol 2019, Isambert and Didier 2024).

Even though temporary fences are effective at reducing roadkill, they do not always prevent individuals from reaching the road (Smith and Sutherland 2014, Gould et al. 2024). Beaune (2019) observed that over 10% of the migrating amphibians were killed on the road despite the installation of a temporary fence, a rate incompatible with the persistence of small populations (Gibbs and Shriver 2005, Beaune 2019). The individuals could have bypassed the fence by its extremities, climbed over it, or crossed under the fence if it was not properly buried. The former behaviour has been witnessed and is known to generate excess mortality at fence extremities (Huijser et al. 2016, Helldin and Petrovan 2019). It can be dealt with by curving the fence in a U-turn (Brehme et al. 2021) or by placing a pitfall trap or a tunnel at each extremity of the fence (Helldin and Petrovan 2019). Mortality linked to individuals climbing over the fence would indicate an improper fence design that should be improved, but knowledge is lacking on this subject. Temporary fence design and materials vary greatly between installations. Wire netting (meshing size 6.5 x 6.5 mm) and polyethylene netting fences are commonly used but amphibians can climb over them in the absence of a sufficiently large overhang (Arntzen et al. 1995, Conan et al. 2022, Gould et al. 2024) . Polyethylene tarp fences have been shown to be effective at reducing trespass rates (Woltz et al. 2008) but were not tested in comparison with other materials. To our knowledge, other temporary fence studies are only descriptive and have not tested the fence effectiveness in controlled conditions (Ganet and Vacher 2012, Gould et al. 2024).

The lack of comparative information about temporary fence materials and designs prevents their optimisation while effective designs are strongly needed as amphibian populations continue to decline in Europe (Helldin and Petrovan 2019, Chanson et al. 2023). This study thus focused on comparing commonly used temporary fences (wire meshing, polyethylene netting and polyethylene tarp) in controlled conditions. We extended the methodology used in Conan et al. (2023) to test higher fences and temporary fence materials. We could then compare materials providing toeholds (wire meshing and polyethylene netting) with a smooth material (polyethylene tarp), and attempt at finding a fence height that agile frogs cannot jump over (Conan et al. 2023). Our aim was (1) to determine the most effective fence configuration, defined as the combination of material, height, and overhang that prevents the highest proportion of amphibians from crossing the fence. We expected long distance jumpers to cross higher fences, and we expected tall and smooth materials to be more effective at blocking amphibians. Our aim was also (2) to find the most efficient temporary fence, defined as the configuration maximising cost-effectiveness, durability, reparability, and ease of use, in order to make recommendations tailored to the capabilities of the various stakeholders.

## MATERIALS & METHODS

### Model species

Two amphibian species with contrasted adult modes of locomotion were used. The common toad (*Bufo bufo* Linnaeus, 1758) represented species moving by walking or hopping. It has a stout body with short hind limbs, and individuals can grow up to 11 - 18 cm in snout-vent length depending on the latitude (Speybroeck et al. 2018). The agile frog (*Rana dalmatina* Fitzinger in Bonaparte, 1839) represented species mainly moving by jumping. It has a smaller frame, long hind limbs and can grow up to 8 cm (Speybroeck et al. 2018). Both species are known to climb on fences. The maximum jumping distance of toads of the genus *Bufo* has been measured in the 10-30 cm range while species of the genus *Rana* were in the 30-200 cm range (Emerson 1978, Zug 1985, Speybroeck 2018).

Forty individuals of each species were captured during their migration season (February 25^th^ 2025 for the common toads and March 22^nd^ 2025 for the agile frogs) and tagged using an RFID PIT. Only males were used because females have been shown to trigger oviposition in similar experiments (Conan et al. 2023). For each species, 30 individuals came from a site equipped with temporary fences (wire meshing) and 10 from a site without fences (designated as ‘naive’) in order to control for a potential effect of learning and memory on crossing performances. Ten individuals were randomly assigned per condition to allow the use of statistical analyses while limiting our impact on wild population as much as possible. Every individual was measured (SVL: from the snout to the end of the ischium) on capture using an electronic calliper (0-150 mm, accuracy: ± 0.03 mm, Tesa technology®, Renens, Switzerland). Captured toads measured from 57.43 to 66.92 mm in SVL (mean 62.66, SE = 0.38) and captured frogs measured from 42.03 to 56.63 mm in SVL (mean 49.48, SE = 0.70). Amphibians were also weighted daily using an electronic scale (Aroma-zone, ± 0.01 g) in order to assess body mass fluctuations during captivity. Capture and housing conditions are detailed in **Supp. 1**. Captures were authorised under the decree AP N° 2024-DREAL-EBP-0054, delivered by the French DREAL Grand Est. The implantation protocol used during this study was validated by the ethics committee for animal experimentation n°036 (CE036; #53332 – 2025012917445217 v 3).

### Experimental setup

All tests were conducted inside a 2000 m^2^ fenced semi-natural outdoor enclosure located between two motorways, Duttlenheim, France (48.512594 °N, 7.582062 °E) in four arenas built out of wood (1*0.5*1 m, length*width*height; **Fig. 1**). The arenas were placed on bricks and underneath a polyethylene tarp to keep them out of the rain and isolate them from the ground. Arenas were adapted from Conan et al. (2023), and consisted of two compartments: (A) ‘departure’ (25x50 cm) which consisted of a movable wooden platform (unattractive) that allowed testing multiple fence heights, while the (B) ‘arrival’ compartment (75x50 cm) was enriched with moist soil, moss and shelters made of branches. Behind the rear side of the ‘arrival’ compartment, a speaker (Pulsar® 2 × 3 W speaker, Enkhuizen, Netherlands) played mating calls from *B. bufo* and *R. dalmatina* to motivate amphibians to cross the fence during the trials as male calls (conspecific or not) are known to attract resident amphibian species (Muller & Schwarzkopf 2017, Testud et al. 2020). The top and the front of the arena were closed with plexiglass, with thin openings on the top to allow for ventilation. The two compartments were separated by a fence. The fence material differed between arenas, with three different materials used: (1) polyethylene tarp (T), (2) 1 mm polyethylene netting (N), and (3) 6.5 mm wire meshing (W, **Fig. 2**).

**Figure 1:**
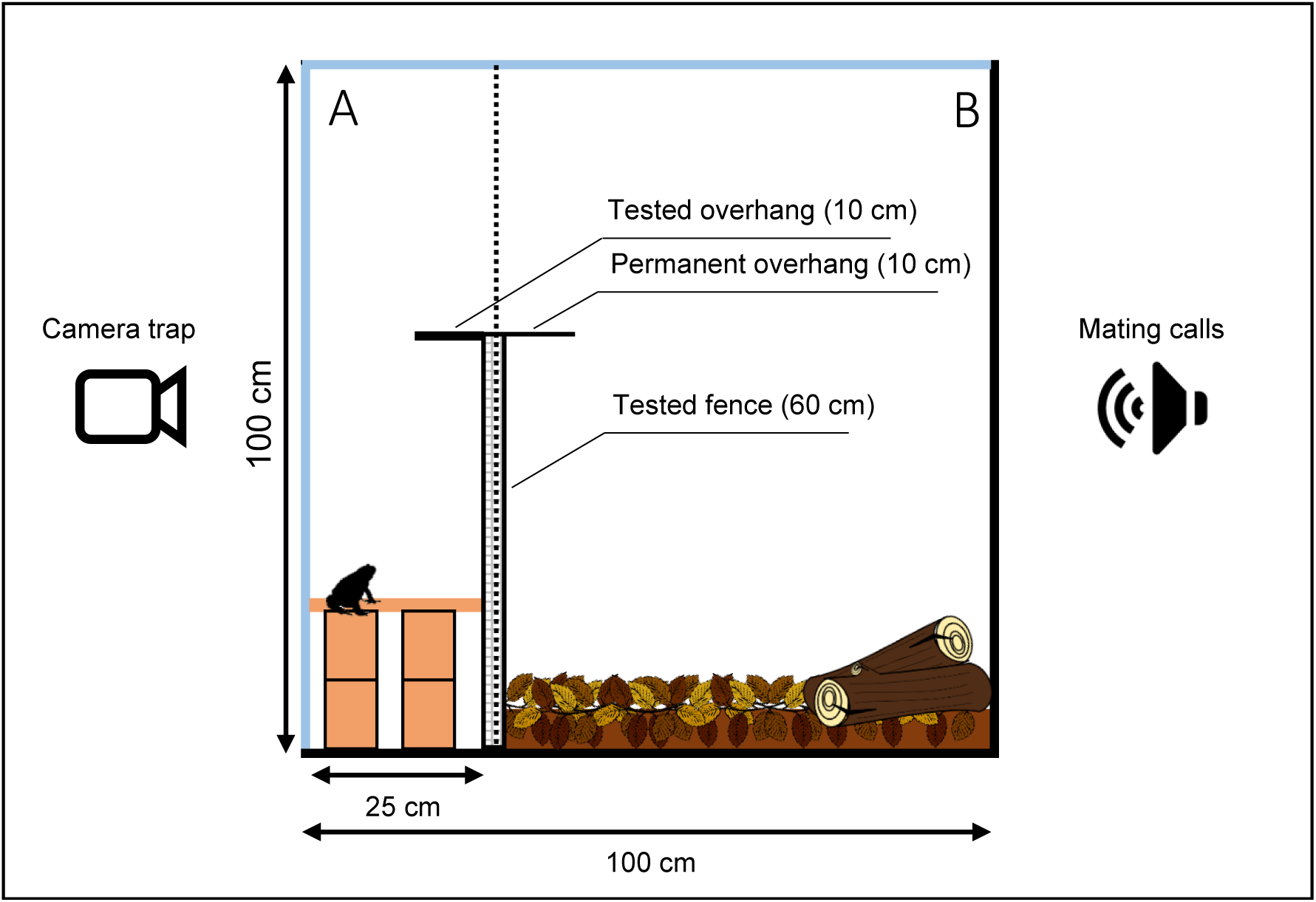
Side view of the experimental arenas. A, the departure compartment composed of a movable wooden floor placed on 10 cm wooden blocks to set up the fence height. B, the arrival compartment composed of moist forest floor and shelters. The tested overhang could be folded back in order to test the condition or not, a permanent overhang prevented individuals from crossing back into the departure compartment. The black sides of the box represent wooden panels; the light blue sides represent plexiglass panels. The Departure compartment was photographed every five minutes by a camera trap; mating calls were played from behind the Arrival compartment to attract amphibians.

**Figure 2:**
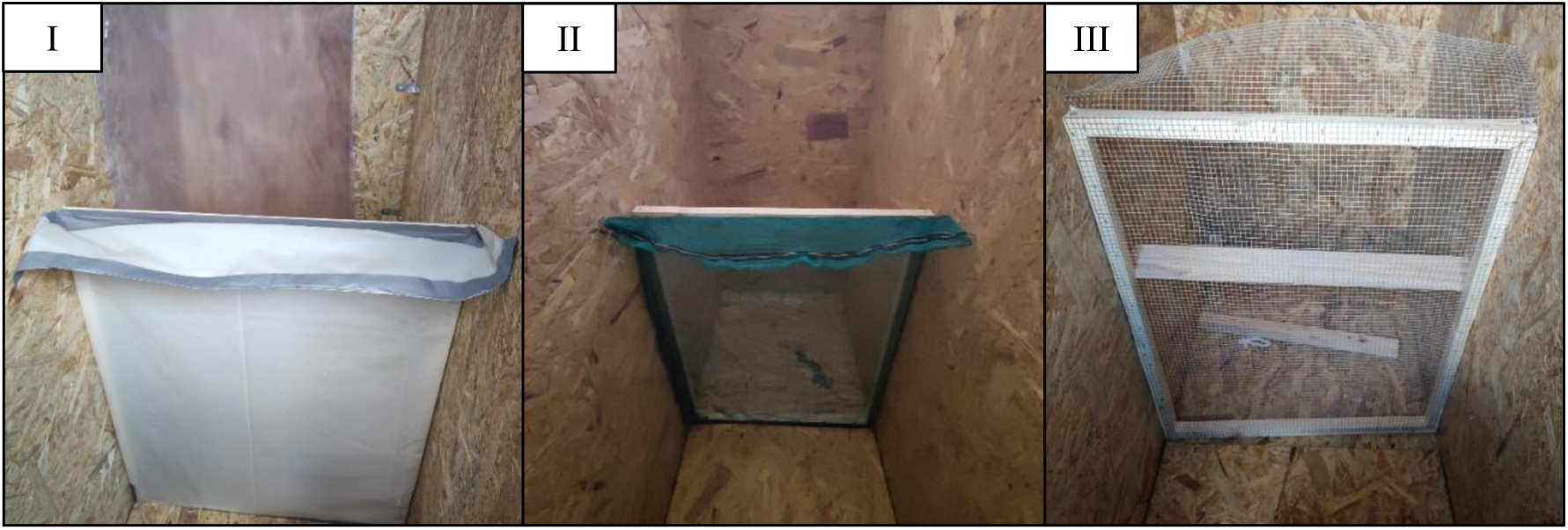
Tested fence materials. I) the polyethylene tarp (T), II) the polyethylene netting (N), and III) the wire meshing (W, Wn). The overhang was set up to be tested in I and II and folded back in III.

Wire meshing fences were used in two arenas: one for the group ‘’wire meshing’’ (W), and one for the group ‘naive’ (Wn). Other materials were not used to test the experience of amphibians because they were not commonly used in the field. Each fence was equipped with a 10 cm horizontal movable overhang to test the impact of the overhang compared to a straight fence. Its dimensions were chosen according to results of previous studies, in which a 10 cm overhang was sufficient to prevent the crossing of most amphibian species (Conan et al. 2022, 2023). The platform of the departure compartment was mounted onto wooden blocks (height = 10 cm) in order to make the fence height vary from 20 to 60 cm with a 10 cm increment.

### Experimental trials

In order to test the three fence materials (polyethylene tarp T, polyethylene netting N, and wire meshing W) and the experience of the amphibians under the W material (naïve Wn or experienced W), the 40 individuals of both species were separated into four groups of 10 individuals. There was one group per fence material (T; N; W) and one for the naive individuals (Wn). The denomination of ‘’group’’ (*e.g.* ‘’tarp group’’) refers hereafter exclusively to the tested individuals, while the denomination of ‘’fence’’ (*e.g.* ‘’tarp fence’’) refers to the physical object. Before starting the trials, each group could explore the arenas with their fence removed, during one habituation night. Given the nocturnal activity of the tested species, all experiments were conducted at night between February 26^th^ 2025 and April 11^th^ 2025, and only on nights for which the minimum forecasted temperature was at least 5°C. At the beginning of a trial, all 10 individuals of each group were placed into the ‘departure’ compartment of their arena and had until the following morning to cross the fence. Trials started at 08:00 PM and ended at 08:00 AM (12 hours), however a logistical constraint made one trial (6^th^ April) last 17 hours for the agile frogs as we had to start the trial earlier than usual. During the entire trial duration, photos were taken every five minutes using camera traps (Cuddeback X-Change™ Color - Model 1279). Camera traps were placed one meter in front of the arena to record amphibian behaviour and interactions with the fence during the night. The following morning, individuals were removed from the arenas and were identified (PIT tags) to determine their crossing status (blocked = 1 /passed = 0). They were then transferred back to their holding tanks.

We tested five fence heights (20, 30, 40, 50 and 60 cm). The 10 cm height was excluded as it was shown to be ineffective for similar species in Conan *et al*., (2023). To minimise the time that amphibians spent in captivity, we tested fence heights and implemented stopping conditions to the testing of fence heights differently for the two species. Toads were expected to be stopped by lower fence heights, so we tested them in the following order: 20, 30, 40, 50, and 60 cm. If every individual were blocked by a given fence height, this height was consecutively tested for up to two additional replication trials. If no individuals crossed the fence during the two replication trials, we considered the fence height as the minimum effective fence height for blocking the amphibians (H_eff_). If some individuals managed to cross the fence during a replication trial, we stopped replicating this fence height and resumed increasing the fence height. The trials were stopped once H_eff_ was found. In contrast, frogs were expected to cross at least 40 cm (Conan et al. 2022, 2023), and easily cross lower fences. Because of that they were tested the fences in the order 60, 50, 40, 30, and 20 cm. If every individual managed to cross the fence during a trial, we did not test lower fences and attributed a crossing success to all individuals for these fences as it meant the frogs were physically able to cross them. Once all fence heights were tested once, replication trials were done starting at the lowest height at which no individuals could cross the fence anymore. Replication trials were done using the same method than for the common toads. These procedures were repeated for both overhang conditions (with and without). When some amphibians ended up crossing the fence during the replication trials of a fence height, we only kept that specific trial for the analyses. When no individuals crossed the fence during the initial trial and the two replication trials of a fence configuration (H_eff_), we only kept the initial trial. It allowed us to have a homogeneous number of data points over fence heights by keeping only one trial per height, while focusing on worse case scenarios, which are the most important to account for in drift fence design.

Night temperatures at the nearby weather station of Holtzheim (∼5 km) were downloaded from the opendatasoft.com website.

### Analyses & statistics

A scan sampling method (Altmann 1974) was used to analyse the behaviour of the individuals with a five-minute time step, corresponding to the delay between each photo (ethogram in **Table 1**).

**Table 1:** Ethogram for the scan sampling behavioural analyses.

| Behaviour | Description |
|---|---|
| Interaction | The individual has at least one limb grabbing or pushing up against the fence |
| Other activity | The individual is active without interacting with the fence |
| Resting | The individual is lying on the ground or on a conspecific |
| Out of sight | The individual is not visible |

Since individuals could not be identified in the photos, the sampling occurred at the group scale, with up to 10 behaviours recorded per scan depending on the number of individuals still present in the departure compartment. For each group, the number of interactions with the fence per night was calculated. For each night and group, the interactions with the fence were recalculated for each hour as the proportion of the Interaction behaviour in overall activity (Overall activity = Interaction + Other activity, **Table 1**). Expressing the interactions with the fence as a proportion of all active behaviours allowed this variable to be independent from the varying number of individuals present in the departure compartment due to fence crossing.

#### Effectiveness

The effect of the group (T, N, W, Wn), height (20, 30, 40, 50 and 60 cm), and their interaction with the overhang (present, absent) was tested on the individual crossing status of agile frogs using ANOVAs based on binomial generalised linear models (**Supp. 2** for general model type and parametrisation). The common toads were tested using the same methodology, but the ‘’overhang’’ variable and the ‘’tarp (T)’’ modality were excluded as they could never cross the fences in these conditions, making the variance for these modalities zero. We considered these modalities significantly different from modalities that allowed some amphibians to cross the fence. The effects of the mean nighttime air temperature, individual SVL, date, and the number of interactions with the fence were included in the model as fixed covariables to help isolate the effect of the fence treatments (Bennett 1984, Swoap et al. 1993, Navas et al. 1999, Oromí et al. 2010, Moreno-Rueda et al. 2020, Conan et al. 2023). The models being binomial, they estimated the probability of the fence blocking the amphibians, which we used to describe fence effectiveness. Effective fence height (H_eff_) was also compared between groups (T, N, W, Wn), but no statistical analyses could be performed as there was only one effective fence height per fence and overhang condition.

#### Efficiency

We defined efficiency in two ways. Firstly, as the interactions with the fence. Fences with which amphibians interacted less were considered more efficient since they would prevent exhaustion. To assess this for agile frogs, we tested the effect of group (T, N, W, Wn), fence height, the presence of the overhang, and time elapsed since the beginning of the trial on the interactions with the fence using ANOVAs based on a beta zero-inflated generalised linear mixed model (**Supp. 2**). Mean night temperature was used as a control variable, and date was included as a random effect. The common toads were tested in the same manner, but only using groups (N, W, Wn) with no overhang to match the nights used for the effectiveness analysis. Because of that, the analysis for the common toads did not require the use of a random variable and the model used was a beta zero-inflated generalised linear model (**Supp. 2**).

Secondly, we explored the cost-benefits balance of each fence concerning its deployment in the field. The following criteria were assessed: cost per 100 meters of fence, mechanical durability, reparability, complexity of installation, and handleability. Cost was the mean price for fences of each material, gathered from readily available online store information and from the suppliers of local operators. Lifespan was either the lifespan or UV resistance lifespan advertised by the manufacturer or the replacement rate used by local operators for the material. Mechanical durability, reparability, installation complexity, and handleability were compared based on the feedback of local operators. Prices and currency conversion rates of December 2025 were used (**Supp. 3**).

Analysis and figures were performed using R software® (R Core Team 2025; v 2024. 12.1+563; packages: car, DHARMa, dplyr, emmeans, ggeffects, ggplot2, glmmTMB, MuMIn, stats) with α = 0.05 significance level. Models underwent variable selection through model averaging (Ullah & Wang, 2013) (**Supp. 2**). Model averaging was done by computing a model selection table based on Bayesian information criterion (BIC) using the dredge function. The average estimators of the global model were then computed by combining the estimators of each sub model in the model selection table, pondered by sub model BIC weight. Average estimators with a significant effect were kept for the final model while other variables were discarded. Tukey HSD post-hoc tests were used when more than two categorical modalities had to be compared. Behavioural sampling was performed with the BORIS® software (v 9.4.1; Friard & Gamba, 2016).

## RESULTS

A total of 27 nights of trials were conducted: 10 nights for toads and 17 nights for frogs. Out of these, only 5 nights were kept to model the probability of blocking toads, and 14 for the probability of blocking frogs after having removed the additional and unsuccessful replication trials used to confirm H_eff_, and after having removed 3 trials during which individuals managed to slip under the wooden floor of the departure compartment. The stopping condition did not affect the homogeneity of the number of trials per fence height used in the analyses for toads because the modalities for which it was applied (tarp fence and fences equipped with an overhang, at any height) were always excluded from the analyses because their variance was equal to zero. The behavioural analyses used the same nights as the fence crossing analyses, except for five trials of the agile frogs. The 20 cm trials of the wire fences without overhangs (W and Wn) were not done due to the stopping condition having been met at 30 cm the day before. The photographs of the 40 cm trial of the netting fence with and without an overhang, and of the 30 cm wire fence without an overhang were not saved on the camera traps due to an unknown issue. From these nights, 3936 single behavioural events were recorded for the common toad and 7009 for the agile frog. Common toads produced a mean of 8.52 ± 0.37 active behaviours per hour (interaction + other activity, **Table. 1**). Agile frogs produced a mean of 8.87 ± 0.30 active behaviours per hour. Towards the end of the trials of the agile frogs, two individuals from the tarp group died and one was lost. The first individual that died could not partake in the last five trials. The second individual that died could not attend the last four trials. Finally, the lost individual could not attend the last two trials.

### Effectiveness

The individual SVL, and number of interactions with the fence had no effect on the probability of the fence blocking the amphibians as they were always discarded during model averaging (**Supp. 2**). The mean night air temperature was also discarded for the common toads but was kept for agile frogs.

#### Probability of blocking the amphibians

When the overhang was present, fences had a moderate to high probability of blocking the agile frogs (**Fig. 3 I**), ranging from 0.67 95CI[0.54 – 0.83] (polyethylene netting, N) to 0.87 95CI[0.71 – 0.95] (naive wire, Wn), with no significant differences between materials (Tukey contrast for N/Wn: odds ratio = 0.27, se = 0.20, z ratio = -1.78, p = 0.63). When no overhang was present, the probability of blocking the agile frogs did not change for the tarp (T) and polyethylene netting (N) fences (**Fig. 3 I**). However, when no overhang was present, the wire meshing fences (W and Wn) had a lower probability of blocking agile frogs than when the overhang was present, at W: 0.33 95CI[0.19 – 0.45] and Wn: 0.30 95CI[0.16 – 0.44] (ANOVA: χ² = 26.57, df = 3, p < 0.01, **Fig. 3 I**). The degradation of the probability of blocking the agile frogs linked to the removal of the overhang amounted to W: - 0.38, and Wn: −0.57, with no significant difference between them. Experience had no effect on the probability of the fence blocking the agile frogs as the naive and non-naive wire groups performed equally (**Fig. 3 I**). Higher fences had a higher probability of blocking the agile frogs (slope = 0.09, se = 0.01, z = 6.46, p < 0.01, **Fig. 3 III**). For agile frogs, the effect of fence height did not differ by material or according to the presence of an overhang either since the interaction of these variables was discarded during model selection (**Supp. 2**). The probability of blocking the agile frogs decreased by 0.06 (se = 0.01) per degree Celsius (slope = −0.49, se = 0.15, z = -3.34, p < 0.01, **Supp. 4**).

**Figure 3:**
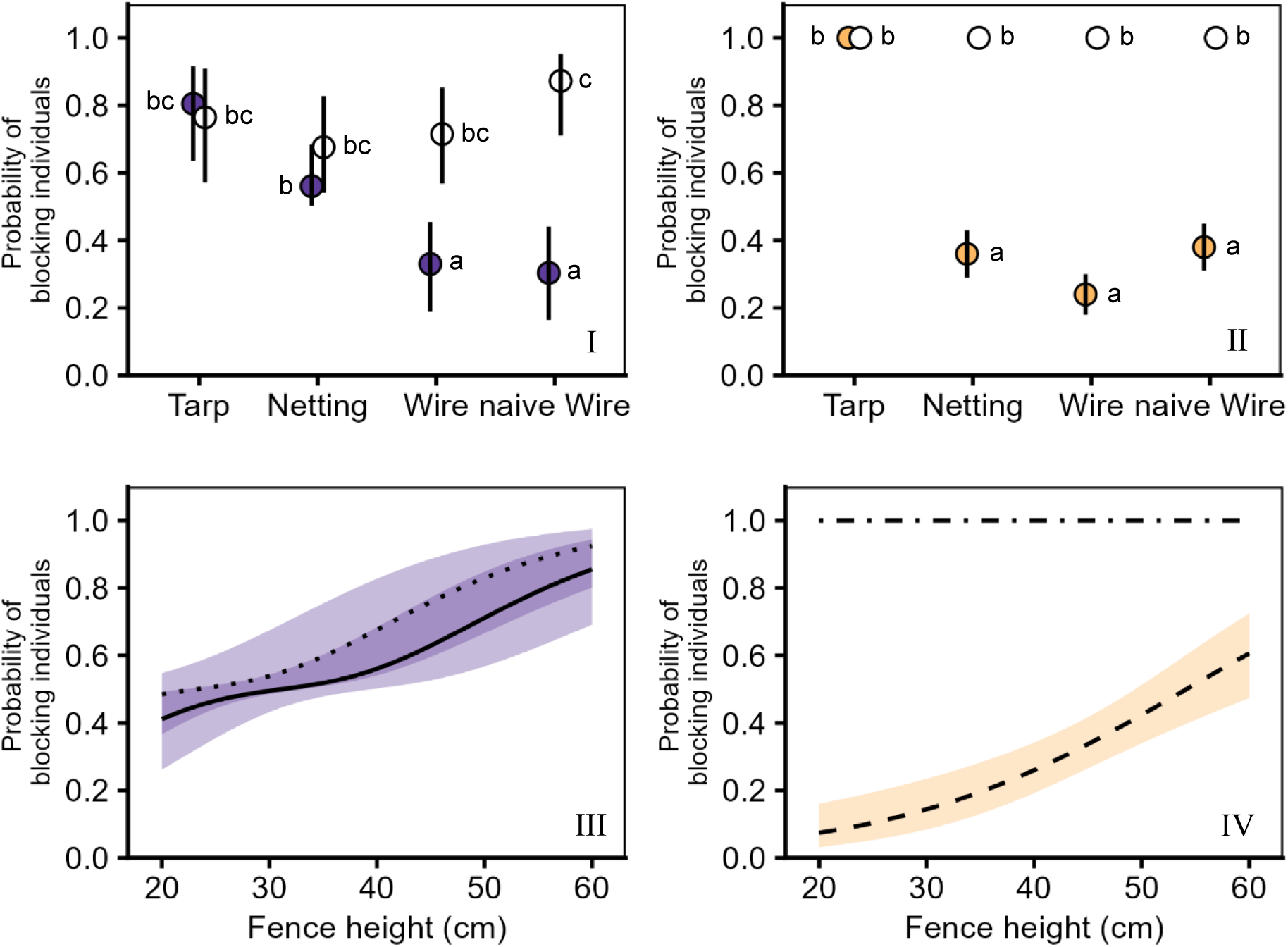
I) Estimated probability of blocking the agile frogs for each fence material, with 95 % confidence intervals. II) Observed probability of blocking the common toads for each fence material, with standard error. Fences with no overhang are represented with coloured, and fences equipped with the overhang are represented with the empty dots. III) Estimated effect of fence height on the probability of blocking the agile frogs, with 95% confidence intervals. The solid line represents fence with no overhang (n = 40 individuals) and the dotted line the fences equipped with an overhang (n = 40 individuals). IV) Estimated effect of fence height on the probability of blocking the common toads, with 95% confidence intervals. The dashed line represents the N, W, and Wn fence with no overhang (n = 30 individuals) and the dot-dashed line the T fence and all fences equipped with an overhang (n = 50 individuals). Frogs are represented by the purple colour and toads by the yellow. Letters a, b, and c represent significant difference levels of the Tukey HSD, with values sharing a letter not differing from each other. The effect of fence height for toads was only calculated based on N, W, and Wn because T always blocked the individuals, thus could not be included as a modality in the model (modality variance = 0).

The common toads were always blocked by the fences when an overhang was present, regardless of fence height or material (**Fig. 3 II, IV**). When the overhang was absent, the tarp fence still blocked every common toad regardless of the fence height (**Fig. 3 II, IV**). When no overhang was present, polyethylene netting (N), and wire meshing fences (W and Wn) had a low probability of blocking the toads, at N: 0.36 (se = 0.07), W: 0.24 (se = 0.06), and Wn: 0.38 (se = 0.07) respectively. There was no significant difference between these materials as the group covariable was discarded during model selection. The degradation in the probability of blocking the toads was −0.64 (N), −0.76 (W), and −0.62 (Wn). Experience had no effect on the probability of the fence blocking the common toads as the naive and non-naive wire groups performed equally (**Fig. 3 II**). The effect of fence height on the probability of blocking toads could only be modeled on the netting (N) and wire (W, Wn) fences with no overhang since the other conditions always stopped them, resulting in their variance being null.

For those materials (N, W, Wn), higher fences had a higher probability of blocking the toads (slope = 0.08, se = 0.02, z = 4.99, p < 0.01), though the probability of blocking the individuals was moderate at best (60 cm: 0.65 [0.51 – 0.78], **Fig. 3 III**). There was no difference between materials (N, W, Wn) since the interaction between these variables was discarded during model selection (**Supp. 2**).

#### Minimum effective height

Polyethylene netting and wire meshing fences were climbable when not equipped with an overhang and were ineffective at any height for both species (**Table 2**). Tarp fences and any fence equipped with an overhang were impossible to climb, they needed to be 60 cm high to prevent agile frogs from jumping above them, while 20 cm were enough to block the common toads (**Table 2**).

**Table 2:** Minimum effective fence height of each fence material, with and without an overhang, and the number of trials it took to identify it for the common toad and the agile frog.

| Fence material | Overhang | Common toad |  | Agile frog |  |
|---|---|---|---|---|---|
|  |  | Minimum effective height | N trials | Minimum effective height | N trials |
| Polyethylene tarp | No | 20 cm | 3 | 60 cm | 9 |
| Polyethylene tarp | Yes | 20 cm | 3 | 50 cm | 5 |
| Polyethylene netting | No | Ineffective | 5 | Ineffective | 6 |
| Polyethylene netting | Yes | 20 cm | 3 | 50 cm | 5 |
| Wire meshing | No | Ineffective | 5 | Ineffective | 1 |
| Wire meshing | Yes | 20 cm | 3 | 60 cm | 8 |

### Energy and cost efficiency

There was no effect of the mean air night temperature, the fence height, and the presence of the overhang on the proportion of interactions with the fence in overall activity for both species as these variables were discarded during model averaging (**Supp. 2**). In addition, time elapsed since the beginning of the trial also had no effect for the common toads, and the group (T, N, W, Wn) had no effect for the frogs as they were also discarded during model averaging (**Supp. 2**).

#### Proportion of interaction with the fence

Common toad individuals tested with the tarp fence interacted with the fence 1.9 times less than the other groups (**Fig. 4 I;** χ² = 22.08, df = 3, p < 0.01). Agile frog interactions with the fence decreased over time during the trials (slope = - 0.12, se = 0.02, p < 0.01, **Fig. 4 II**). Experience had no effect on the interactions with the fence since there was no difference between the naive (Wn) and non-naive wire (W) groups for the common toads (**Fig. 4 I**), and the group variable was discarded during model selection for the agile frogs.

**Figure 4:**
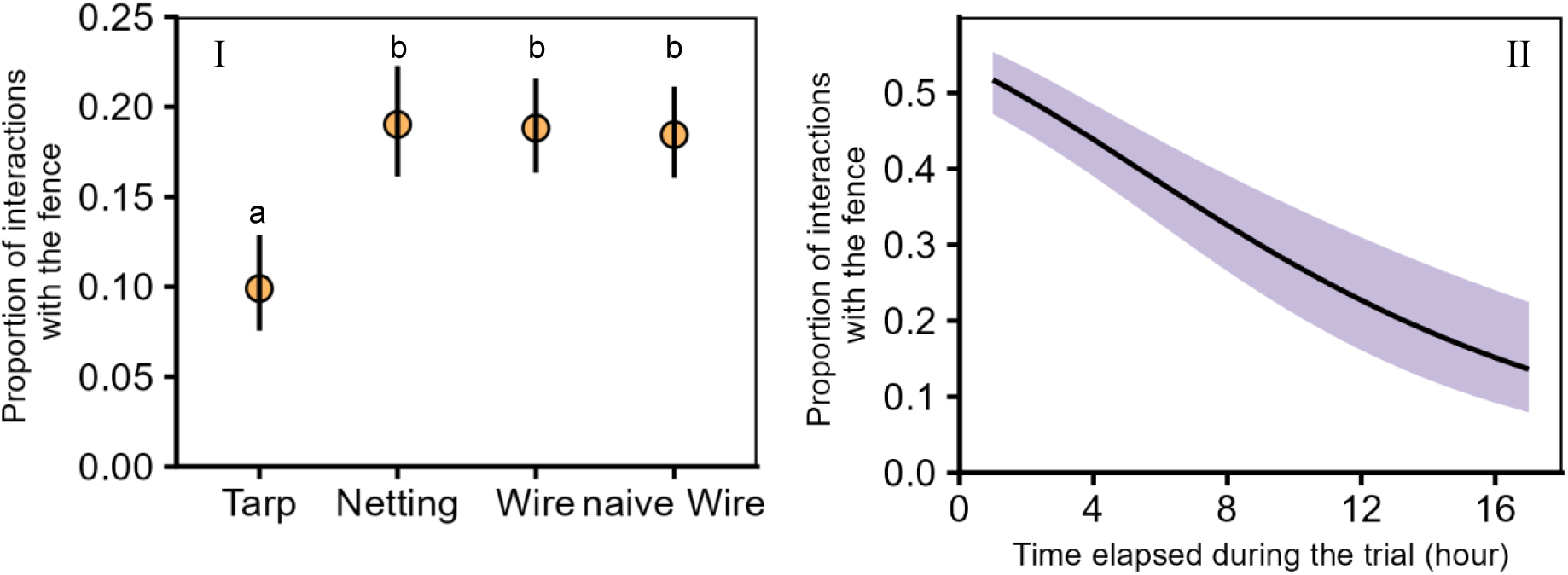
Predicted effect and 95 % confidence intervals of I) fence material and II) time elapsed since the beginning of the trial on the proportion of interaction with the fence in global activity. Common toads are in yellow and agile frogs in purple. Toads interacted less with the tarp fence but kept interacting equally during the night while frogs interacted less with the fence as time passed regardless of the fence material.

#### Cost-benefits

Fourteen references from 10 manufacturers were gathered from online searches for readily available prices combined with a quote for material used by a local operator (**Supp. 3**). The polyethylene netting fences were the most cost-efficient option (**Table 3**). They had the cheapest cost per 100 m per year of lifespan at 54.17 € ± 11.51 and were the easiest to handle as they were light, resistant enough to be manipulated with no extra care, and could not hurt operators. They were technically less resistant than wire meshing fences, but both could be damaged by wild boars leading to a similar functional durability. The tarp fence was quickly and easily repaired as it only needed heavy-duty duct tape to be put back together. The other materials were repaired by sewing the netting or the wires back together, which was a longer task to perform. The polyethylene tarp fence was however the most complex one to set up, requiring bent poles, guiding threads for the overhang, and tension threads to keep the fence upright. The easiest fence to set up was the wire meshing fence which only needed straight poles and a manual bending of its top to create the overhang. The lack of a guiding structure however meant that the overhang could be improperly formed, rendering it ineffective. Eight local operators gave us feedback after having tried installing polyethylene netting instead of their usual wire fences. Two were positive, three neutral, and three negative. The positive feedback reported an easier and faster installation of the polyethylene netting. The negative feedback concerned the increased complexity of the netting fence and that it was harder to set up in sandy grounds.

**Table 3:** Cost/benefits analysis of the three fence types studied.

|  | Polyethylene tarp | Polyethylene netting | Wire meshing |
|---|---|---|---|
| Cost (€/100 m) | 116.71 ± 10.28 | 325.49 ± 69.10 | 895.79 ± 178.61 |
| Lifespan (years) | 2 (UV lifespan) | 6 (UV lifespan) | 10 (replacement rate) |
| Cost/year (€/100m) | 58.35 ± 5.14 | 54.25 ± 11.52 | 89.58 ± 17.86 |
| Mechanical durability | Puncture/tears by mesomammals, blown off by wind/semitrailers | Puncture/tears by macromammals | Puncture/tears by macromammals |
| Reparability | Duct tape | Sewing (polyethylene thread) | Sewing (steel wire) |
| Installation complexity | Curved poles, guiding thread | Curved poles, guiding/ tension threads | Straight poles, fence bent to form the overhang |
| Handleability | Risk of puncture | No drawback | Heavier, wire can cause hand injuries |

## DISCUSSION

We showed that the most effective fences configuration at blocking the amphibians are the tarp fence, or any fence equipped with an overhang. When having one of these configurations, 20 cm fences managed to block common toads, while it took 60 cm fences to block agile frogs. We also showed that polyethylene tarp and polyethylene netting fences had a similar cost per year, with the tarp being easier the repair and less complex to install, at the cost of durability and handleability.

### Experimental design & limitations

We tried to reproduce realistic conditions during the trials by placing the arenas outdoors, close to a motorway, as temporary fences are usually installed next to noisy areas like roads or construction sites. We also tried to maximize the motivation of the individuals by placing them at night, during the migration season, on a non-attractive substrate after having allowed them to learn the existence of a suitable substrate before the trials (Dall’antonia and Sinsch 2001). Arenas were set side-by-side in a 1 by 2 meters rectangle in a position ensuring a homogeneous exposition to external disturbances. We were able to isolate the effect of our treatments on the crossing success and activity properly by accounting for the effect of temperature and body size, and the cumulated number of interactions with the fence during the trials. Temperature is one of the most important drivers of ectotherm muscle activity (Navas et al. 1999, Arnfield et al. 2012, Gunderson and Leal 2015), and body size can directly influence locomotor abilities (Cayuela et al. 2020, Moreno-Rueda et al. 2020). In our case, body size likely had no effect due to the low variation in the SVL of the tested individuals. The effect of fatigue over the different trials was accounted for by using the cumulated number of interactions with the fence during each trial as a covariable as exhausted amphibians would have been less active. This variable was always discarded during model selection, meaning that exhaustion should not have affected our results.

Two technical points are to be taken in account for the arenas to function properly in the setup we used. The wooden floor of the ‘’departure compartment’’ should be tightly fitting with the walls of the arena and the fence on all four sides, otherwise amphibians will manage to slip under the floor and not complete the trial. Duct taping any gap left after setting up the arena is an easy way to make the wooden floor frog-proof. The second point of attention is condensation: the plexiglass will rapidly be covered with fog if the arena is not well ventilated, obscuring the individuals from the camera traps. It can be solved by drilling small holes in the roof of the arena or by integrating mesh in it.

Despite our precautions, our experimental design cannot eliminate every bias linked to testing groups of individuals, as interactions between individuals may have differed between groups and affected the outcome of the experiment. Previous experience from scan and focal sampling analyses of similar fence crossing trials (Conan et al. 2023) had shown no interactions between individuals other than resting amphibians sometimes lying on top of each other (unpublished data). It allowed climbing individuals to reach the top of smooth fences up to 13 cm. The use of 20 cm as a minimum fence height in this study should have avoided this effect. If any bias of this nature remains in our results, it should only affect the comparison between treatment groups (fence materials) and not the effect of fence height or the presence of the overhang within groups.

### Effectiveness

#### Walker and jumper species

Fences presenting toeholds (wire meshing and polyethylene netting) always allowed at least one individual to cross them when no overhang was present. However, minimum fence height was found in conditions designed to prevent the amphibians from crossing the fence by climbing (polyethylene tarp or overhang present). This difference indicates that these anti-climbing designs are indeed effective at blocking the amphibians. It means that in non-climbable conditions (tarp fence or overhang present), the amphibians could only cross the fences by jumping above them. The 20 cm and 60 cm minimum fence heights of the toads and frogs respectively indicate that these species cannot jump. For conditions in which amphibians could cross the fences, higher fences blocked a higher proportion of the individuals for both species, although it was less effective for toads than for frogs. This suggests either that toads are better climbers than frogs, or that when climbing is an option, it is a more reliable method for crossing fences compared to jumping.

In order for fences to be effective for walkers and long-distance jumpers, their height needs to be adapted to the most agile species present in the area of deployment, and their structure should prevent the possibility to climb over the fence via the use of an overhang or smooth materials. Our findings extend previous results on fence crossing in amphibians (Conan et al. 2022, 2023) to the most common species found crossing roads in Europe (*B. bufo*) and highlight that even a mesh size of 1 mm can be used as a toehold. We also completed the results from Conan et al. (2023) on the agile frogs by finding them unable to jump above 60 cm fences, which converges with results on *L. pipiens* (Woltz et al. 2008). The agile frog has arguably the best jumping abilities in Europe so this result can be used as higher limit for fence design in the region. From our findings and the available literature, 60 cm high fences seem to be a minimum to ensure over 90 % of effectiveness for long-distance jumpers.

#### Adherence climber species

Adherence climbers were initially supposed to be included in this study with the Palmate newt (*Lissotriton helveticus* Razoumovsky, 1789) but an unusually low numbers of migrating individuals prevented us from capturing enough of them to include in our protocol. Even though testing them would have provided more accurate data, we can infer from previous studies that newts should be able to climb all three materials tested in our study as long as no overhang is present and if the fences are wet (Morand & Carsignol 2019, Boissinot *et al*. 2023, Csutoros 2025). Adhesive toe pads are a common trait in frog morphology (Barnes et al. 2011) making tree frog species proficient climbers on almost any surface (Hertwig and Sinsch 1995, Federle et al. 2006, Crawford et al. 2016, Langowski et al. 2018). Given the existing literature on tree frogs, we would expect them to easily climb any material and only be stopped by a 40 cm+ fence equipped with an overhang (Dodd 1991, Conan *et al. 2023,* Gould *et al*. 2024).

### Energy and cost efficiency

The tarp was the most effective material, but our results show that the type of material used for the fence is not critical as long as a 10 cm overhang is present, which we strongly advise for in any cases since it always improves the effectiveness of the fence. The addition of an overhang allows to shift the focus of the choice of material from fence effectiveness to efficiency and cost-benefits, a core part of mitigation fences design (Huijser et al. 2009, 2022).

#### Wire meshing

From our results, the wire meshing was the least efficient of the three types of materials. Not only is the cost per year of these fences twice as much as the other two materials, but smaller newts (*Lissotriton sp.*) can get stuck in the meshing or crawl through it (Association Les Piverts 2024, and our personal observations). The mean skull width of adult *Lissotriton helveticus* (∼7 mm; Bettencourt- Amarante et al., 2024) is smaller than the diagonal of a 6.50 mm square (9.19 mm), the commonly used mesh size in France (Morand & Carsignol 2019, Conan et al. 2022), which allows newts to pass their heads through it. A mesh size of 3 mm (4.24 mm diagonal) should prevent this. The advantages of wire meshing do not counterbalance its cost since it can still be damaged by wildlife and might require maintenance during the migration period, and the overhang folding can be done incorrectly on large portions of fence (Jumeau 2017).

#### Polyethylene tarp

The two polyethylene materials were similar in term of cost/durability. The tarp was more fragile as it could be easily punctured and blown off by wind gusts but was more easily repaired than the netting, making it more suited for short term projects. Our results showed reduced interaction of toads with the tarp fence, which could be linked to its opacity as shown in chapters 4, 5, and 6 of Brehme et al. (2020). This phenomenon could be beneficial if it prevents toads from exhausting themselves trying to climb the fence but could be detrimental if it ends up hindering the guiding function of drift fences.

#### Polyethylene netting

The longer longevity of polyethylene netting fences makes them more suited for repeated use. A potential drawback of the netting could be an abrasion-linked trauma that was witnessed on the snout of frogs near polyethylene netting fences by Gould *et al*. (2024), although they did not witness the trauma happening. We did not observe such trauma during our trials but if the netting was proven to cause such abrasion, the material should then be avoided. Polyethylene netting was the most complex fence to set up. When compared to the least complex fence (wire meshing), 63% of operators were satisfied with this material or found it easier to set up, while 37% found it more difficult to set up. Accounting for the fact that it was the first time the operators used the polyethylene fence, the added complexity of this design might not be a major drawback in the end, especially for experienced operators.

## CONCLUSION

In a context where temporary drift fences for amphibians still allow for some trespass in the field, we managed to identify the most effective fence material for amphibians, which was the polyethylene tarp as it was the only one able to prevent crossings without an overhang. Materials presenting toeholds can still make effective amphibian fences on the condition that a 10 cm overhang is present at the top of the fence, the addition of an overhang being mandatory regardless of materials if adherence climbers are present or suspected on site. Regardless of the material used, fence height should be at least 60 cm to ensure high effectiveness for long-distance jumper species. A cost-benefits comparison of these effective designs highlighted two designs depending on the duration of the fence deployment. For repeated use, we recommend 60 cm polyethylene netting fences with a 10 cm, 90° overhang as they are the best combination of effectiveness, cost and durability. For punctual use and if wind/ truck traffic is not a problem, we recommend 60 cm tarp fences with a 10 cm 90° overhang since this option is three times cheaper than the polyethylene netting.

## ACKNOWLEDGEMENTS

This study was funded by the French Collectivité européenne d’Alsace. We thank Mila Witcher for their work in helping to build the experimental arenas, running the experimental trials and sampling behaviours from pictures, and every other student that occasionally helped with running the experimental trials. We also thank the Ligue pour la Protection des Oiseaux Alsace association and the Eurométropole d’Alsace for their collaboration. Finally, we thank the Massif forestier de Strasbourg Neuhof – Illkirch Graffenstaden nature reserve, the Office National des Forêts, as well as Mr & Mrs Moge and Mr Walther for allowing us access to the breeding ponds present on their properties.

## AUTHORS CONTRIBUTION

Meven Le Brishoual, Claude Miaud and Jonathan Jumeau conceived the study and designed the experimental protocols. Meven Le Brishoual conceived the experimental arenas and built them with Nathan Dehaut. Meven Le Brishoual, Clara Tanvier and Nathan Dehaut collected the data in the field. Meven Le Brishoual and Clara Tanvier analysed the data. Meven Le Brishoual and Clara Tanvier drafted the manuscript. Jonathan Jumeau, Claude Miaud and Nathan Dehaut gave critical feedback on the manuscript. All authors have read and approved this version of the manuscript for publication and agree to be held accountable for their contributed part of the work.

## DATA AVAILABILITY

All of the data and R scripts used for the statistical analyses are available online from the Zenodo repository https://doi.org/10.5281/zenodo.24159051 (Le Brishoual et al. 2026).

All supplementary materials are available on the bioRxiv preprint server at https://www.biorxiv.org/content/10.64898/2026.04.14.718167v8.supplementary-material

## CONFLICT OF INTEREST

The authors declare no conflicts of interest.

## Notes

https://doi.org/10.5281/zenodo.24159051

### Competing Interest Statement

The authors have declared no competing interest.

### Summary of Updates

This version follows the 4th round of revision of the article with mainly changes in formulations to improve clarity. The figure about the effect of temperature on the probability of blocking the agile frogs has been moved to the supplementary materials (Supp. 4).

