## Supplementary material for "Effectiveness and efficiency of temporary fences as mitigation measures for preventing amphibian roadkill": Supp. 1

### Supplementary 1.1 Capture

All individuals were captured at their breeding site during the breeding season (February-March 2025). Captures were conducted manually using dip nets during nocturnal patrols. To prevent contamination by pathogens between sites, nitrile gloves were used and the wet gear was disinfected using Virkon S 1 % (Lanxess; Miaud, 2022). For each species 40 individuals were captured, including 30 individuals from a site equipped with a temporary fence (‘‘non-naive’’ groups, common toads: 49.497462, 7.609378, common frogs: 48.835459, 7.676666), and 10 from another site without one (‘‘naive group’’, common toads: 48.552032, 7.502045, common frogs: 48.702933, 7.721281). Distinction between naive and non-naive groups was used to control for a potential effect of learning and memory on crossing performances. Only adult males were used in the experiment in order to reduce the impact on wild populations since female anurans can lay eggs under stress (Conan et al. 2023). Only adult individuals were tested because drift fences are usually no longer present in the field when juveniles are ready to cross roads (Schmidt and Zumbach 2008). Moreover, previous studies already covered wire meshing and slick fences for juveniles (Conan et al. 2022, 2023). To ensure individual tracking during experimentation, individuals were implanted subcutaneously and on the left flank with a RFID PIT tag (1.4×9 mm; TAG LF GLT1M4X9 RO EM, Biolog-ID®, Bernay, France), under local anaesthesia, at the time of capture. Two morphological measurements were taken: (1) body mass in g using an electronic scale (Aroma-zone, ± 0.01 g), and (2) snout-vent length in mm (SVL: from the snout to the end of the ischium) was measured using an electronic calliper (0–150 mm, accuracy: ±0.03 mm, Tesa technology®, Renens, Switzerland).

### Supplementary 1.2 Housing

Experimental arenas and housing were located inside a 2000 m^2^ fenced semi-natural outdoor enclosure, Duttlenheim, France (48.512594 °N, 7.582062 °W). Between trials, individuals were kept indoors in 800×600×435 mm polypropylene tanks with perforated lids (Euronorm container 12.4046.07 AXESS industries®, Strasbourg, France). Individuals were housed together by species with up to 10 individuals per tank for common toads and up to 20 for agile frogs. Tanks were enriched with moist soil; flat tiles and stones provided shelters for the individuals. Water and food were provided *ab libitum* in the tanks. Water was provided in cups (for soaking), and food consisted of crickets (*Acheta domesticus*, ~1 cm) and Wax moth larvae (*Galleria mellonella*, ~2 cm). Temperature was not controlled in the tanks and was similar to the external temperature. Body mass was recorded every morning following nightly trials to monitor animal health. Individuals showing abnormal weight loss were excluded from tests until recovery. Experiments could last for a maximum of 20 days, after which all individuals were released at their respective capture locations.
