## Supplementary material for "Effectiveness and efficiency of temporary fences as mitigation measures for preventing amphibian roadkill": Supp. 2

| *Supplementary 2.1: Model parametrisation of the general models used for the model averaging procedure with species concerned, model types (and families), response variables, fixed variables, random variables, random variable variances, and detection of zero inflation.* | | | | | | |
| --- | --- | --- | --- | --- | --- | --- |
| Species | Model type | response variable | fixed variables | random variables | rand. variance | Zero inflation |
| *B. bufo* | GLM  (binomial) | individual crossing status | group * height + SVL + temperature + interactions w/ fence | - | - | no |
| *B. bufo* | ZIGLM  (beta) | interactions with the fence | group + height + temperature + time | - | - | yes |
| *R. dalmatina* | GLMM  (binomial) | individual crossing status | group * overhang * height + SVL + temperature + interactions w/ fence | individual | 0.10 | no |
| *R. dalmatina* | ZIGLMM  (beta) | interactions with the fence | group * overhang + height + temperature + time | date | 0.05 | yes |
| ***individual crossing status****: whether the individuals has been blocked (1) or not (0)*  ***interactions with the fence****: proportion of interactions with the fence in total active behaviours*  ***group****: material test group the individual (T, N, W, Wn)*  ***overhang****: presence of the overhang at the top of the fence (yes/no)*  ***height****: fence height for the trail (20, 30, 40, 50, 60)*  ***SVL****: snout-vent length of the individual in cm*  ***temperature****: mean air temperature during the night of the trial*  ***interactions w/ fence****: cumulated number of interactions with the fence for the group during the night of the trial*  ***time****: number of hours elapsed since the beginning of the trial*  ***date****: date of the trial*  ***individual****: PIT number of the individual* | | | | | | |

| *Supplementary 2.2: Model averaging table for the individual crossing status of common toads with general model variable, model estimate, estimate standard error, adjusted standard error, z-value, and p-value* | | | | | | |
| --- | --- | --- | --- | --- | --- | --- |
| Variable | estimate | se | adjusted se | z-value | p-value |  |
| Intercept | -8.33 | 4.22 | 4.24 | 1.97 | <0.05 | * |
| Group (W) | -0.03 | 0.18 | 0.18 | 0.15 | 0.88 |  |
| Group (Wn) | <0.001 | 0.11 | 0.11 | 0.04 | 0.97 |  |
| Fence height | 0.14 | 0.04 | 0.04 | 3.57 | <0.001 | *** |
| Fence height : group (W) | <0.001 | <0.01 | <0.01 | 0.15 | 1.00 |  |
| Fence height : group (Wn) | <0.001 | <0.01 | <0.01 | 0.04 | 0.99 |  |
| SVL | -0.01 | 0.05 | 0.05 | 0.31 | 0.76 |  |
| Temperature | 0.47 | 0.25 | 0.25 | 1.90 | 0.06 |  |
| Interaction | -1.00e-3 | <0.01 | <0.01 | 0.2 | 0.84 |  |

| *Supplementary 2.3: Model averaging table for the individual crossing status of agile frogs with variable, model estimate, estimate standard error, adjusted standard error, z-value, and p-value* | | | | | | |
| --- | --- | --- | --- | --- | --- | --- |
| Variable | estimate | se | adjusted se | z-value | p-value |  |
| Intercept | 2.90 | 2.52 | 2.52 | 1.15 | 0.25 |  |
| Group (T) | 1.50 | 0.62 | 0.62 | 2.40 | 0.02 | * |
| Group (W) | -2.37 | 0.61 | 0.61 | 3.89 | <0.001 | *** |
| Group (Wn) | -2.51 | 0.62 | 0.62 | 4.06 | <0.001 | *** |
| Overhang (present) | 0.76 | 0.68 | 0.69 | 1.11 | 0.27 |  |
| Group (T) : overhang (present) | -0.97 | 0.91 | 0.91 | 1.06 | 0.29 |  |
| Group (W) : overhang (present) | 2.57 | 0.87 | 0.87 | 2.95 | <0.01 | ** |
| Group (Wn) : overhang (present) | 3.76 | 0.94 | 0.97 | 3.77 | <0.001 | *** |
| Fence height | 0.09 | 0.02 | 0.02 | 6.14 | <0.001 | *** |
| Fence height : overhang (present) | 7.51e-4 | 7.46e-3 | 7.48e-3 | 0.10 | 0.92 |  |
| Fence height : group (T) | 2.06e-5 | 1.27e-3 | 1.28e-3 | 0.02 | 0.99 |  |
| Fence height : group (W) | 3.42e-5 | 1.70e-3 | 1.71e-3 | 0.02 | 0.98 |  |
| Fence height : group (Wn) | -6.82e-7 | 9.11e-4 | 9.13e-4 | 1.00e-3 | 1.00 |  |
| Fence height : overhang (present) : group (T) | 2.74e-9 | 2.43e-5 | 2.44e-5 | 0.00 | 1.00 |  |
| Fence height : overhang (present) : group (W) | -5.58e-9 | 3.05e-5 | 3.06e-5 | 0.00 | 1.00 |  |
| Fence height : overhang (present) : group (Wn) | -7.14e-9 | 3.59e-5 | 3.59e-5 | 0.00 | 1.00 |  |
| SVL | -0.01 | 0.03 | 0.03 | 0.37 | 0.71 |  |
| Temperature | -0.46 | 0.18 | 0.18 | 2.51 | 0.01 | * |
| Interaction | 8.81e-4 | 5.11e-3 | 5.12e-3 | 0.17 | 0.86 |  |

| *Supplementary 2.4: Model averaging table for the interactions with the fence of common toads with variable, model estimate, estimate standard error, adjusted standard error, z-value, and p-value.* | | | | | | |
| --- | --- | --- | --- | --- | --- | --- |
| Variable | estimate | se | adjusted se | z-value | p-value |  |
| Zero-inflation | -0.34 | 0.13 | 0.13 | 2.54 | <0.05 | * |
| Intercept | -1.534 | 0.15 | 0.16 | 8.66 | <0.001 | *** |
| Group (T) | -0.71 | 0.20 | 0.20 | 3.574 | <0.001 | *** |
| Group (W) | 0.01 | 0.13 | 0.13 | 0.08 | 0.94 |  |
| Group (Wn) | -0.01 | 0.13 | 0.13 | 0.04 | 0.97 |  |
| Fence height | -1.22e-4 | 1.19e-3 | 1.19e-3 | 0.10 | 0.92 |  |
| Temperature | 2.02e-3 | 0.01 | 0.01 | 0.19 | 0.85 |  |
| Time | -0.03 | 0.02 | 0.02 | 1.23 | 0.22 |  |

| *Supplementary 2.5: Model averaging table for the interactions with the fence of agile frogs with variable, model estimate, estimate standard error, adjusted standard error, z-value, and p-value.* | | | | | | |
| --- | --- | --- | --- | --- | --- | --- |
| Variable | estimate | se | adjusted se | z-value | p-value |  |
| Zero-inflation | 0.38 | 0.10 | 0.10 | 3.74 | <0.001 | *** |
| Intercept | -1.33 | 0.38 | 0.38 | 3.52 | <0.001 | *** |
| Group (T) | -0.17 | 0.23 | 0.23 | 0.74 | 0.46 |  |
| Group (W) | -0.02 | 0.08 | 0.09 | 0.22 | 0.83 |  |
| Group (Wn) | -0.01 | 0.09 | 0.09 | 0.07 | 0.95 |  |
| Overhang (present) | -0.01 | 0.07 | 0.07 | 0.16 | 0.87 |  |
| Group (T) : overhang (present) | -6.31e-5 | 0.01 | 0.01 | 0.01 | 0.99 |  |
| Group (W) : overhang (present) | 3.26e-4 | 0.01 | 0.01 | 0.02 | 0.98 |  |
| Group (Wn) : overhang (present) | 5.78e-4 | 0.02 | 0.02 | 0.03 | 0.98 |  |
| Fence height | 2.13e-4 | 2.01e-3 | 2.02e-3 | 0.11 | 0.92 |  |
| Temperature | 4.96e-3 | 0.03 | 0.03 | 0.18 | 0.86 |  |
| Time | -0.13 | 0.02 | 0.02 | 5.95 | <0.001 | *** |
