## Supplementary material for "Effectiveness and efficiency of temporary fences as mitigation measures for preventing amphibian roadkill": Supp. 3

| *Supplementary 3: Comparison of online readily available amphibian temporary drift fences with fence model name, company name, fence material, price per unit, length of one unit in meters, price per 100 meters (converted in euros when needed, November 2025), and UV resistance lifespan. For UV resistance, DD: lack of information; Yes: branded as UV resistance with no lifespan indicated; NA: no sensitivity to UV.* | | | | | | |
| --- | --- | --- | --- | --- | --- | --- |
| Produce name | company | material | price | length (m) | price/100m € | UV resistance |
| Ecofender standard | Hy-Tex UK ldt | Polyethylene tarp | 108.00 £ | 100 | 123.12 | 2 years |
| Caudon temporary newt fencing (clear) | Legacy Habitat management ldt | Polyethylene tarp | 112.85 £ | 100 | 128.65 | DD |
| Caudon temporary newt fencing (green) | Legacy Habitat management ldt | Polyethylene tarp | 113.11 £ | 100 | 128.95 | DD |
| Caudon NewtX temporary newt fencing | Legacy Habitat management ldt | Polyethylene tarp | 75.54 £ | 100 | 86.12 | DD |
| toad fence amphibian safety fence 200g/m² | Dr. Thiel manufacturing textile solutions | Polyethylene netting | 389.00 € | 100 | 389.00 | 6 years |
| toad protection fence | Grube KG | Polyethylene netting | 375.00 € | 100 | 375.00 | Yes |
| amphibian protection fence | Grube KG | Polyethylene fabric | 499.00 € | 95 | 525.26 | Yes |
| HDPE amphibian protection | Hermann Meyer KG | Polyethylene netting | 178.90 € | 100 | 178.90 | Yes |
| filet B. Vent F1132 | Jost SA | Polyethylene netting | 159.30 € | 100 | 159.30 | DD |
| small welded wire mesh sheets - 6 mm | Origin Suregreen ldt | Wire mesh | 9.00 £ | 0.9 | 1140.00 | NA |
| 6mm x 6mm Stainless steel wire mesh  (H90cm x L30m) – 22g/0.7mm | Wire Fence | Wire mesh | 240.29 £ | 30 | 913.10 | NA |
| grillage métallique électrosoudé 1 m x 5 m  FENSANET 6,4 | Nortene Home Depot France | Wire mesh | 57.99 € | 5 | 1159.80 | NA |
| Grillage volière 6.5x6.5x0M50 de haut, fils de 0.7 mm d’épaisseur | SARL Euro-Negoces | Wire mesh | 50.68 | 25 | 202.72 | NA |
| grillage pour animaux soudé gris, H.1 x L.3 m,  maille H.6 x l.6.4 mm Centrale brico | Vianal Management SAS | Wire mesh | 31.90 € | 3 | 1063.33 | NA |
