## Supplementary material for "Effectiveness and efficiency of temporary fences as mitigation measures for preventing amphibian roadkill": Supp. 4

*Supplementary 4: Estimated effect of the mean air temperature during the trials on the probability of blocking the agile frogs, with 95 % confidence intervals. The fences blocked the agile frogs less at hotter temperatures.*

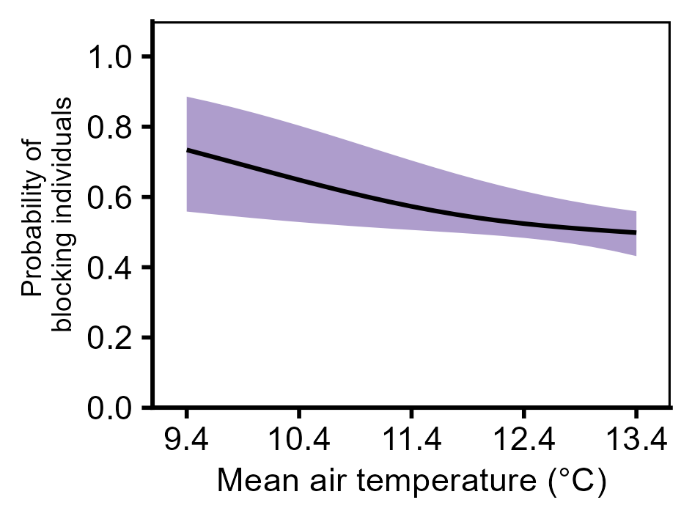
